# Time-dependent modulation of spinal excitability during action observation in the macaque monkey

**DOI:** 10.1101/768804

**Authors:** S.J. Jerjian, R.N. Lemon, A. Kraskov

## Abstract

Neurons in the primate motor cortex, including identified pyramidal tract neurons projecting to the spinal cord, respond to the observation of others’ actions, yet this does not cause movement in the observer. Here, we investigated changes in spinal excitability during action observation by monitoring short latency electromyographic responses produced by single shocks delivered directly to the pyramidal tract. Responses in hand and digit muscles were recorded from two adult rhesus macaques while they performed, observed or withheld reach-to-grasp and hold actions. We found modest grasp-specific facilitation of hand muscle responses during hand shaping for grasp, which persisted when the grasp was predictable but obscured from the monkey’s vision. We also found evidence of a more general inhibition before observed movement onset, and the size of this inhibition effect was comparable to the inhibition after an explicit NoGo signal. These results confirm that the spinal circuitry controlling hand muscles is modulated during action observation, and this may be driven by internal representations of actions. The relatively modest changes in spinal excitability during observation suggest net corticospinal outflow exerts only minor, sub-threshold changes on hand motoneuron pools, thereby preventing any overflow of mirror activity into overt movement.

---

Mirror neurons are found in the ventral premotor cortex (PMv), primary motor cortex (M1) and other parts of the primate motor cortical network. These cells modulate their firing during both the execution of grasping actions and the observation of similar actions performed by others (Gallese et al. 1996; Rizzolatti et al. 1996; Kilner and Lemon 2013). Activity in this latter condition occurs in the absence of any detectable muscle activity or movement in the observer, despite the finding that even pyramidal tract neurons (PTNs), which project directly to the spinal cord, can be mirror neurons (Kraskov et al. 2009; Vigneswaran et al. 2013). The extent to which cortical mirror activity influences the excitability of spinal motor networks is not well understood.

PMv, where mirror neurons were first discovered, has limited direct projections to the lower cervical spinal cord (He et al. 1993; Borra et al. 2010), but mirror neurons in PMv could also exert influence at the spinal level via strong anatomical (Muakkassa and Strick 1979; Godschalk et al. 1984; Matelli et al. 1986; Dum and Strick 2005) and functional (Cerri et al. 2003; Shimazu et al. 2004; Schmidlin et al. 2008; Prabhu et al. 2009; Kraskov et al. 2011; Maier et al. 2013) connectivity with M1. M1 itself makes a major contribution to corticospinal projections to the cervical cord, including via its direct corticomotoneuronal (CM) projections to hand muscle motoneurons (Porter and Lemon 1993; Rathelot and Strick 2006; Lemon 2008). Importantly, many mirror PTNs, particularly in M1, either show sharply reduced activity during action observation compared with execution, or actually suppress their activity relative to baseline (Vigneswaran et al. 2013). Together, these features could feasibly produce a “cancellation effect” to prevent mirror activity from inducing movement (Kraskov et al. 2009, 2014; Vigneswaran et al. 2013). However, it is difficult to predict the spinal impact of cortical PTN activity (or lack of it) without knowing their targets and actions, which could include both excitation and inhibition at the spinal level (Soteropoulos 2018).

In humans, transcranial magnetic stimulation (TMS) has been useful for non-invasively probing net excitability changes during action observation. TMS over M1 during observation of grasp generally results in an increased amplitude of motor evoked potentials (MEPs) in hand and arm muscles (Fadiga et al. 1995; Gangitano et al. 2001; Urgesi et al. 2006; Cattaneo et al. 2009), consistent with the congruence of mirror neuron responses during action execution and observation. Short-intracortical inhibition paradigms have suggested that excitability changes during action observation are primarily cortical in origin (Strafella and Paus 2000). Monitoring of H-reflexes in upper limb muscles provides a measure of spinal excitability changes, and several studies have reported fluctuations in H-reflex amplitude during action observation with a profile similar to that of muscle activity occurring during execution of the same actions (Baldissera et al. 2001; Borroni et al. 2005; Montagna et al. 2005). H-reflexes may be confounded by pre-synaptic inhibition, and are difficult to elicit in intrinsic hand muscles (Mazzocchio et al. 1995; Knikou 2008), which are of particular relevance for grasping by virtue of their strong CM connections (Porter and Lemon 1993; McKiernan et al. 1998). The net balance of cortical and spinal effects underlying excitability changes during action observation is uncertain due to variability across subjects and tasks (Naish et al. 2014; Hannah et al. 2018).

Stimulation of the pyramidal tract (PT) produces antidromic responses which can be used to identify PTNs, and also produces orthodromic descending volleys which exert direct and indirect actions on spinal motoneurons. PT stimulation has advantages for probing spinal excitability compared with stimulating the cortex or peripheral afferents, as the evoked descending volley is unaffected by cortical interactions, and is not thought to be subject to the spinal pre-synaptic inhibition affecting peripheral afferent inputs (Jackson et al. 2006). Direct, monosynaptic excitation of hand muscle motoneurons from this volley gives rise to a short-latency MEP in these muscles (Olivier et al 2001; Cerri et al 2003), and the amplitude of the MEP should reflect post-synaptic excitability of the motoneurons during different phases of action observation, including any descending effects originating from mirror neurons. A more complete understanding of the pattern and extent of action observation activity not just within cortical networks, but also in the downstream spinal circuitry controlling hand muscles, could provide new insights into mechanisms for the generation and suppression of grasping movements.

Here, we assessed changes in excitability during action observation in the primate spinal cord via direct stimulation of the pyramidal tract (PT) in two rhesus macaques and comparison of the modulation of short-latency evoked responses in hand muscles. Mirror neurons have previously been shown to continue to modulate when the grasp is obscured (Umiltá et al. 2001), and also been implicated in the suppression of movement (Kraskov et al. 2009, 2014; Vigneswaran et al. 2013) and inaction representation (Bonini et al. 2014). We therefore also examined excitability changes when the availability of visual information of the observed grasp was altered, or an explicit stop signal was provided in place of the implied movement suppression required during passive observation.

## MATERIALS & METHODS

### Animals

Experiments were performed on two adult male purpose-bred macaque monkeys (Macaca mulatta: M48, 12.2 kg and M49, 10.5 kg). All experimental procedures were approved by the UCL Animal Welfare and Ethical Review Body and carried out in accordance with the UK Animals (Scientific Procedures) Act. The monkeys were singly-housed according to veterinary advice, in a unit with other rhesus monkeys, with natural light and access to an exercise pen and forage area. Both monkeys are continuing to participate in experiments.

### Task

The monkey and a human experimenter sat opposite each other, with a custom-designed experimental box placed between them (Fig. 1A). Two homepads faced the monkey, with an additional one in front of the experimenter. Two target objects were present in the peri-personal space of both the monkey and the experimenter - a trapezoid object (base: 30mm height: 27.5mm, width/depth: 20mm) affording precision grip (PG) and a sphere (radius 20mm) affording whole-hand grasp (WHG). The distance between the objects’ centres of rotation was 76mm. Four LEDs surrounded the objects and were used to provide trial information. The monkeys were able to see the objects through a controllable LCD screen (14×10cm), which remained opaque between trials. Trials were initiated after a short inter-trial interval (1-2s), after the monkey had depressed the two homepads with both hands, and the experimenter depressed their homepad on the opposite side. The LCD screen became transparent (LCDon, Fig. 1B), and the object field was illuminated. After a short delay (0.25-0.45s), two orange LEDs indicated the upcoming object and required grasp (PG or WHG) and acted as a warning signal (object cue: ObjCue). After a further delay (0.8-1.2s), the trial type was indicated as execution (green LED on monkey side), observation (green LED on experimenter side), or NoGo (red LED on monkey side). On execution trials, the green LED acted as the Go cue for the monkey to release the right homepad (homepad release; HPR), reach towards, and grasp the target object (displacement onset; DO) using their right hand and the appropriate grasp (Fig. 1A inset). The monkey then clockwise-rotated the object by at least 30°, and held it there for 1s (hold on to hold off; HO-HOFF). The force level required to maintain a grasp was approximately 3N for both objects. A static tone indicated that the monkey was in the hold position, and a second, higher, tone followed to indicate successful completion of the hold. The monkey released the object and returned to the homepad (homepad return; HPN). On observation trials, the roles were reversed, and the experimenter performed the same reach-to-grasp action using their left hand, while the monkey remained still on the homepads.

**Fig 1.**
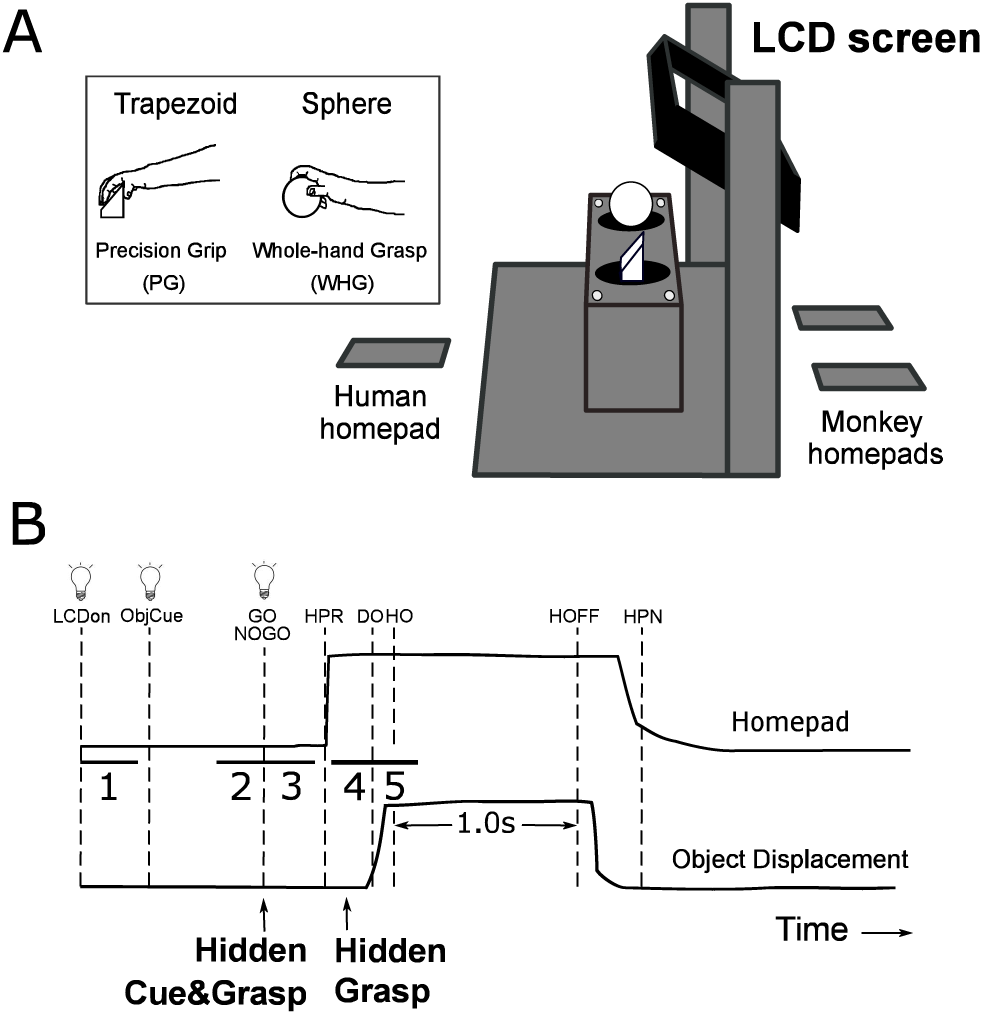
Experimental Task Design. (**A**) The monkey and human experimenter sat opposite each other with the experimental box between them, and rested hands on homepad sensors. The monkey viewed two target objects through a controllable LCD screen. LEDs around the target objects first indicated the upcoming object (ObjCue) followed by the trial type (Execution, Observation, or NoGo). 2 LEDs on experimenter’s side were used on Hidden Cue&Grasp trials to selectively withhold trial information from the monkey. Inset shows the trapezoid and sphere objects, and the precision and whole-hand grasps, respectively, used by the monkey to grasp the objects on execution trials. (**B**) Schematic of object displacement and homepad signals, and corresponding task events on Go trials. Two arrows denote times at which the screen became opaque on hidden trials - Hidden Cue&Grasp (at Go cue), Hidden Grasp (0.15s after experimenter HPR). On Hidden Cue&Grasp trials, object and go cues were not visible to the monkey. Numbered intervals 1-5 indicate epochs used for statistical analysis of MEPs. Vertical lines and labels denote time of task events, detected online by experimental software and used for later alignment. LCDon: LCD screen becomes transparent, ObjCue: two orange LEDs indicate the upcoming object and required grasp; HPR: homepad release, DO: object displacement onset, HO: hold on, HOFF: hold off, HPN: homepad return.

Observation trials were split into three sub-types. On Visible Observation trials, the screen remained transparent until HPN. On Hidden Grasp trials, the screen became opaque 0.15s after HPR, so that the monkey knew the upcoming grasp (via the ObjCue and Go cue), and saw the initial part of the experimenter reach, but could not see the grasp itself. On Hidden Cue&Grasp trials, object cues were provided to the experimenter via different LEDs (not visible to the monkey), and the screen became opaque at the moment of the Go cue. The monkey therefore could not see the grasp, as well as having no information about which object would be grasped. The hold and reward tones were retained for all trial types. On NoGo trials, the red LED instructed the monkey to remain still on the homepads for 1s until a tone indicated successful completion of the trial. The monkey received a small fruit reward for each successfully completed trial of any condition. All trial types and objects were presented in pseudo-random order, in a 20:6:6:3:5 ratio (Execution, Visible Observation, Hidden Grasp, Hidden Cue&Grasp, NoGo), with the higher proportion of Execution trials used to encourage monkeys to remain engaged in the task. Incorrectly performed trials, defined by inappropriate release of a homepad or a violation of timing constraints within the task, were detected and aborted online by the experimental software, not rewarded, and excluded from all analyses.

### Surgeries

Three separate and well-spaced surgical procedures were required to prepare the monkeys for these experiments. First, monkeys were implanted with a ring-shaped TekaPEEK headpiece, secured to the skull for stable head fixation, both for these recordings and other related experiments. In a second surgery, EMG electrodes were subcutaneously tunnelled from a connector on the headpiece through to the right arm and chronically implanted into 12 muscles in the arm and hand. In a third surgery, two tungsten electrodes were stereotaxically implanted in the left medullary pyramid in an anterior-posterior configuration. Each electrode was lowered in 0.5mm steps while single test stimuli of 300μA were delivered at regular intervals. The stimulus configuration was either monopolar against ground on skin, or bipolar between the electrodes. The final electrode position was determined on the basis of the threshold of the antidromic volley recorded via a ball electrode placed on the dura over the left primary motor cortex. Electrodes were fixed at a location eliciting the lowest threshold antidromic volleys (<50μA), with no additional signs of facial or hypoglossal activation (Kraskov et al. 2009). All surgeries were performed under aseptic conditions, under full general anaesthesia (ketamine 0.1ml/kg i/m, maintained with 1.5-2% isoflurane in O_2_). Monkeys were recovered in a padded cage overnight, and treated with antibiotic and analgesic medication before and after surgery.

### PT stimulation

We delivered single bipolar, biphasic stimulus pulses (each phase 0.2ms) between the two chronically implanted electrodes continuously during task performance (300μA every 500ms in M48, 350μA every 600ms in M49). PT stimulation elicited MEPs in the right hand. Stimulus current was monitored at all times, and evoked EMG responses at rest, when present, were of moderate amplitude and submaximal. These parameters were similar to those used in separate sessions for antidromic identification of pyramidal tract neurons in PMv and M1. The stimulus frequency reflected a compromise between obtaining a sufficient number of responses within each session, avoiding any summation across consecutive responses, and ensuring that the monkey tolerated the stimulation and maintained good task performance.

### Recording

Stimulation was delivered to the left medullary pyramid during task performance over 5 separate sessions in each monkey. Monkeys performed roughly 25 trials each in the Visible and Hidden Grasp observation conditions, 12 trials in the Hidden Cue&Grasp observation condition, 20 NoGo trials, and 80 execution trials, per session and object. We simultaneously recorded EMG activity from muscles in the right arm and hand, including three intrinsic hand muscles (first dorsal interosseous (1DI), thenar eminence, and abductor digiti minimi (AbDM)). To confirm absence of EMG activity during action observation, we also recorded from the biceps muscle, and several forearm extensors and flexors (abductor pollicis brevis (AbPL), extensor digitorum communis (EDC), extensor carpi ulnaris (ECU), flexor digitorum profundus (FDP), flexor carpi ulnaris (FCU), flexor carpi radialis (FCR), brachioradialis (BRR)). We also recorded from the extensor carpi radialis longus (ECR-L) in M48, and the deltoid in M49. EMG was hardware high-pass filtered at 30Hz, amplified 2000x or 5000x, and sampled at 5 kHz. We simultaneously recorded the timing of all task events and PT stimuli (25 kHz), and analog object displacement and homepad pressure signals (5 kHz). All signals were stored on an offline computer for later analysis. In separate sessions, we recorded eye movements from each monkey’s right eye using a non-invasive ISCAN camera system (ETL-200, 125Hz), which was located ∼25cm away from, and ∼17cm above the objects. No explicit fixation criteria were imposed on the monkeys during the task.

### Data Analysis

#### EMG

For visualization of overall EMG activity during task performance without PT stimulation, signals were rectified, and bandpass-filtered (30-500Hz, 4th order, zero-phase Butterworth). EMG signals were downsampled to 500Hz, further smoothed with a 100ms moving average and aligned to the Go/NoGo cue for each correct trial, normalized to the 99th percentile across all trials within each channel, and then averaged across trials.

#### Behaviour

Object displacement and homepad signals were also aligned to the Go/NoGo cue for each correct trial, normalized to the 99th percentile across all trials and channels (displacement, monkey homepads or human homepad separately), and averaged across trials. On execution and observation trials, we defined reaction time as the time between the Go cue (LEDon) and HPR, and movement time as the interval between HPR and DO.

### Eye Movements

The location of each object was estimated from a smoothed (24ms moving average) x-y map of eye movements on monkey trials. First, maximally visited locations in the 400ms intervals (+/- 200ms) around Go and HPR were identified. Monkeys typically fixated on the target object LED around the Go signal, and around the object at HPR. The range of horizontal and vertical eye positions visited at least half as often as these maximally visited locations were then used to form a rectangular window estimate for each object’s location. These windows included the objects and their corresponding LEDs on the monkey side, and subtended a visual angle of approximately 5°. The two windows were confirmed to be non-overlapping in the x-direction for each session, but given the unequal position of the two objects relative to the camera, were allowed to have different sizes in the y-direction. Heat maps of gaze behaviour produced for each session clearly indicated well-separated locations of the relevant LED and object during the ObjCue and execution reach-to-grasp periods, respectively.

Within a session, we aligned data to different task events and defined, for each condition and timepoint separately, a dwell likelihood, the proportion of trials in which the monkey’s gaze fell within either the window for the target object, the other object, or neither window. A dwell likelihood equal to 1 for a given 24ms time window thus indicates that the monkey’s gaze fell within the corresponding spatial window at that time window on every trial, whereas a dwell likelihood of 0 indicates that the monkey never looked within the window at that time point. The resulting dwell likelihoods were then averaged across sessions, and expressed as mean±SEM at each time point.

### Motor-evoked potentials (MEPs)

To quantify MEP amplitude modulation across the task, we initially grouped stimuli times across each session into 300ms-long bins aligned to salient task events. 1. −300 to +300ms around LCDon (2 bins), 2. −1200 to +300ms around LEDon (5 bins) 3. −300 to +300ms around HPR (2 bins) 4. −300 to 1500ms around DO (6 bins). Responses to stimuli falling outside any of these bins (e.g. in the inter-trial interval), and the corresponding evoked responses, were excluded from further analysis. For the NoGo condition, only the first 2 alignments (7 bins) were defined. As we recorded EMG in a range of hand and arm muscles, we were able to quantitatively exclude MEPs which may have been influenced by voluntary EMG. To do so, we used the absolute average of all raw pre-stimulus EMG segments (100ms period immediately before the stimulus) in the two bins around LCDon (−300ms to +300ms) to determine muscle-specific thresholds. Any stimulus during passive periods of the task (all observation and NoGo bins, execution bins before the Go cue) for which the average absolute pre-stimulus EMG in any muscle exceeded 5 standard deviations (S.D.) of the mean were discarded. If any trial contained more than 2 contaminated MEPs, all MEPs from that trial were discarded. This resulted in 2.3±0.3% (mean±standard error) of MEPs being discarded in M48, and 1.6±0.8% in M49, with no more than 4% discarded in any session. A stricter threshold of 3S.D. of the mean resulted in higher discard rates (M48: 7.8±0.8% M49: 7.6±0.1%), but did not qualitatively change any of the results. For each muscle and MEP, the peak-to-peak (peak-to-trough) amplitude was extracted from the raw unrectified EMG traces in the 5-14ms period after stimulus delivery. Peak-to-peak amplitudes were then averaged within each bin and session, and these averages were normalized to the average amplitude across conditions of the 300ms bin beginning at LCDon.

### Statistical Analysis

To assess modulation during observation, we performed planned comparisons of MEP amplitude across five epochs of interest for Visible and Hidden Grasp observation conditions via 2-way ANOVA (factors EPOCH x OBJECT (PG, WHG). The five epochs were as follows: 1. *Baseline* (0 to 300ms from LCDon), 2. *ObjCue* (−300 to 0ms before Go/NoGo), 3. *Reaction* (0 to 300ms from Go/NoGo cue), 4. *Hand Shaping* (−300 to 0ms before DO), and *Grasp* (0 to 300ms from DO). The Hidden Cue&Grasp condition was identical for both objects and contained a lower number of trials per object, and showed no significant differences in the 2-way ANOVA. We therefore pooled MEPs from trials for both objects in this condition and assessed modulation via a 1-way ANOVA across the same five epochs.

We performed an additional 2-way ANOVA (factors CONDITION x OBJECT) to compare directly across the 3 observation conditions and 2 objects during the *Hand Shaping* epoch.

To assess modulation of MEPs during NoGo in relation to execution and observation, we compared 2 epochs (ObjCue and Reaction) across 3 conditions (Execution, Visible Observation, and NoGo) and 2 objects via 3-way ANOVA (EPOCH x CONDITION x OBJECT). For all analyses, significant ANOVA results were followed by post-hoc tests where appropriate, with Tukey-Kramer correction for multiple comparisons. P-values ≤ 0.05 (corrected) were considered significant.

All data processing and analysis was performed using custom-written scripts in Matlab 2016b (The MathWorks, Inc., Natick, MA, USA).

## RESULTS

To assess modulation of spinal excitability during action observation, we recorded MEPs elicited by direct stimulation of the medullary pyramid while monkeys passively observed a human experimenter performing cued reach-grasp-and-hold movements. These observation trials were randomly interleaved with execution trials, in which the monkeys performed the same movements themselves, and NoGo trials, where the monkeys were explicitly cued to refrain from movement. Furthermore, on a subset of observation trials, a controllable LCD screen was used to modulate visual information provided to the monkeys about the upcoming grasp.

### EMG and Task performance

EMG activity recorded during sessions without stimulation showed distinct patterns for the different grasps during action execution. In the action observation (full visibility) and NoGo conditions, there was no systematic EMG activity (Fig. 2A and B, note EMG during observation and NoGo is plotted at x10 higher gain). During stimulation sessions, any stimulus-aligned sweeps contaminated by voluntary EMG activity in non-movement stages of the task were removed before analysis (see Methods). Monkey reaction and movement times across all trials were considerably faster than the human experimenter (see supplementary table 1; all monkey-human comparisons p < 0.0001, Mann-Whitney U-test). Monkey reach times were generally longer for PG than WHG, due to the object’s location contralateral to the reaching (right) arm (supplementary table 1, both within-monkey comparisons p < 0.0001, Mann-Whitney U-test).

**Fig 2.**
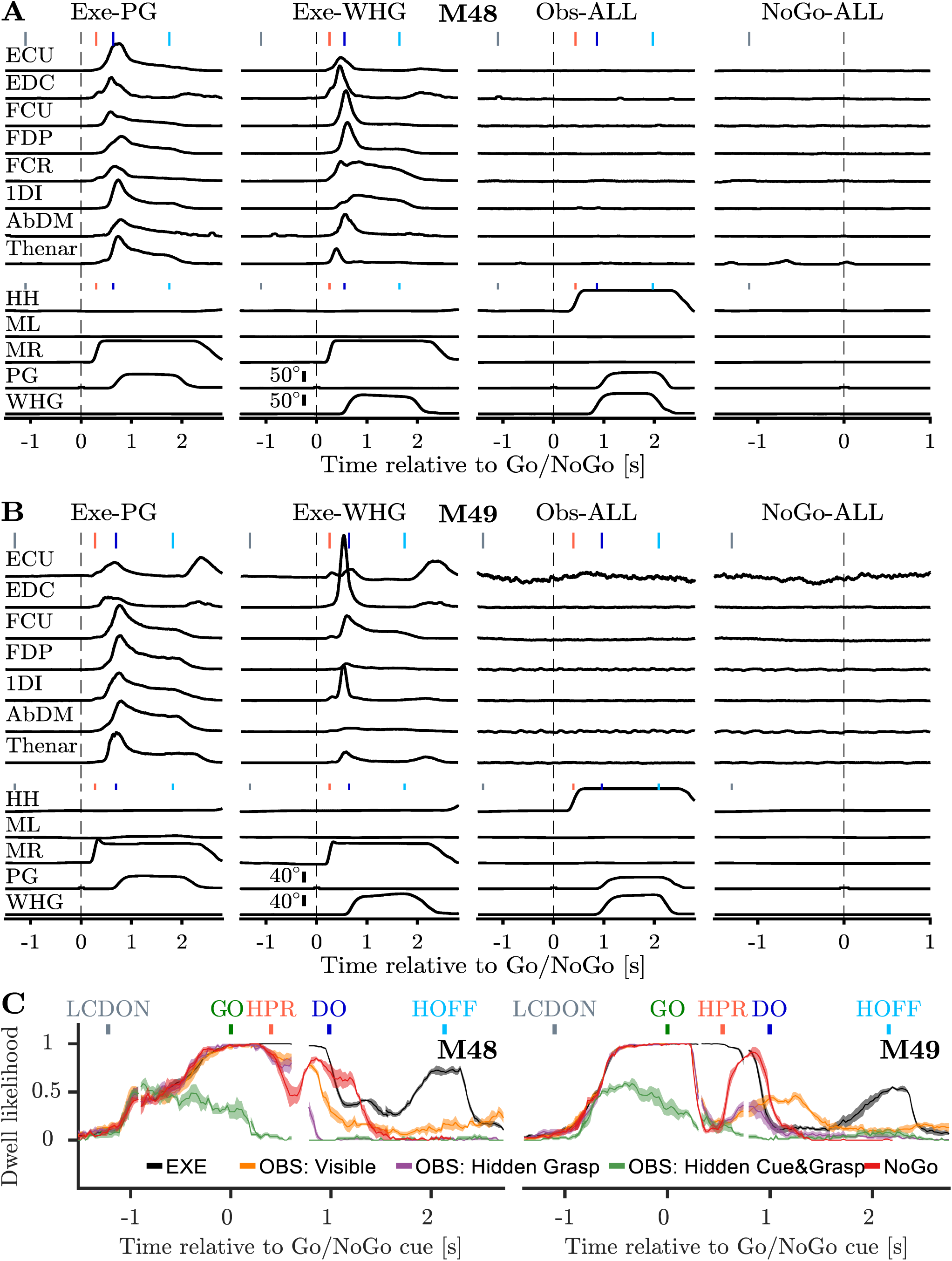
(**A**) Task-related EMG and behaviour from single sessions (without stimulation) in M48. Top panels show pre-processed, rectified, and normalized EMG activity for different muscles (see Methods). Corresponding object displacement and homepad signals are shown in bottom panels. Execution EMG is presented for both objects separately, observation (Visible only) and NoGo EMG are pooled across objects, and shown with a 10x higher gain. Vertical markers at top of each trace indicate median time of key task events relative to Go/NoGo cue (dashed lines). Grey, LCDon: LCD screen becomes transparent; Black (dashed), LEDon, a green (Go trials) or red LED (NoGo trial) for a Go/NoGo signal; Red: HPR: homepad release, Blue, DO: object displacement onset, Light blue HOFF: hold off. (**B**) Same as (A), but for M49. PG: precision grip, WHG: whole-hand grasp, EDC: extensor digitorum communis, ECU: extensor carpi ulnaris, FDP: flexor digitorum profundus, FCR: flexor carpi radialis, 1DI: first dorsal interosseous, AbDM: abductor digiti minimi, HH: human homepad, ML: monkey left homepad, MR: monkey right homepad, PG: trapezoid displacement, WHG: sphere displacement. (**C**) Dwell frequency of gaze on target object in M48 (left) and M49 (right). Traces + flanking shaded regions show proportion of trials for each condition (mean±SEM across 4-5 sessions) in which monkey’s gaze fell into a window around the target object at each timepoint, and for each condition. Data is multiply aligned to different task events (LCDon, Go/NoGo, HPR, DO, and HOFF, colour conventions as in Fig. 2A).

### Eye Movements

In separate sessions, we recorded eye movements in each monkey to assess whether monkeys attended to the observation task, and verify that the opaque screen successfully abolished the visual information available to the monkey about the upcoming grasp. For each trial and time point we defined whether the monkey’s gaze fell within a window around (a) the target object for the given trial, (b) the other (non-target) object, or (c) outside of both windows (see Methods). Fig. 2C shows, for each condition, time point, and monkey, the likelihood that gaze fell within the target object window. A value of 1 indicates that the monkey’s gaze was within the target object window at that particular time point on every trial. These values are averaged across sessions, so that the error reflects the variability across sessions, and the mean value itself reflects variability in gaze within a session. Both monkeys reliably began to look at the target object over the course of the Object Cue period, except during Hidden Cue&Grasp trials where they did not exceed chance (0.5), and looked away after the Go cue (when the screen went off). During Visible and HiddenGrasp observation, M48 maintained gaze until shortly after experimenter HPR, and tended to remain longer and occasionally view the actual grasp and hold on Visible Observation trials. In these conditions, M49 looked away earlier, before occasionally returning gaze to the target object during the reach and grasp.

On execution trials, monkeys maintained gaze until DO (M48) or late reach (M49), frequently looked away while maintaining stable hold, and tended to return gaze to the object in the lead-up to the end of the hold. In the NoGo condition, both monkeys gaze behaviour showed a similar pattern to observation trials, viewing the target object during the ObjCue period and then deviating shortly after the NoGo cue. Both monkeys also returned to the target object towards the end of the trial (1s after the NoGo cue), possibly in expectation of the extinguishing of the LED marking the end of the trial and subsequent reward.

### Motor-evoked potentials (MEPs)

Overall, each stimulation session lasted 30-40 minutes, resulting in 3000-4500 PT stimuli being delivered. Stimuli were binned in 300ms epochs according to their timing relative to task events and sweeps contaminated by pre-stimulus EMG were rejected (<4% of binned MEPs per session, see Methods). Within the epochs used for statistical analysis, there remained an average of 13.5 ± 0.4 MEPs per epoch, object and observation/NoGo condition in M48, and 11.2 ± 0.2 in M49 (MEP counts for each condition and epoch separately are presented in Supplementary Table 2).

Figure 3A shows single sweep responses to PT stimulation in the 1DI muscle (M48) during the observation grasp interval, with a latency of 9-10ms, consistent with monosynaptic activation of spinal motoneurons (Olivier et al. 2001; Cerri et al. 2003). In several muscles in M49, we observed some unusually long-latency effects during passive periods of the task, which sometimes dominated, but were also often present in tandem with short-latency responses (Fig. 3B, see Discussion). For the purposes of this study, we focused on the short-latency responses typical of direct (monosynaptic) excitation of spinal motoneurons. As we did not attempt to optimise the stimulation intensity for each muscle, the amplitude of MEP responses was also variable across muscles. Figure 3C and D show examples of averaged MEPs from five muscles in each subject. For further analysis, we concentrated on distal muscles (1DI, thenar complex and AbDM) most relevant for the grasp, due to the known stronger CM connections to these muscles (Nakajima et al. 2000) compared with proximal muscles (Porter and Lemon 1993; McKiernan et al. 1998; Morecraft et al. 2013). The AbDM muscle in M48 was excluded because of a high level of noise in the recorded signal.

**Fig 3.**
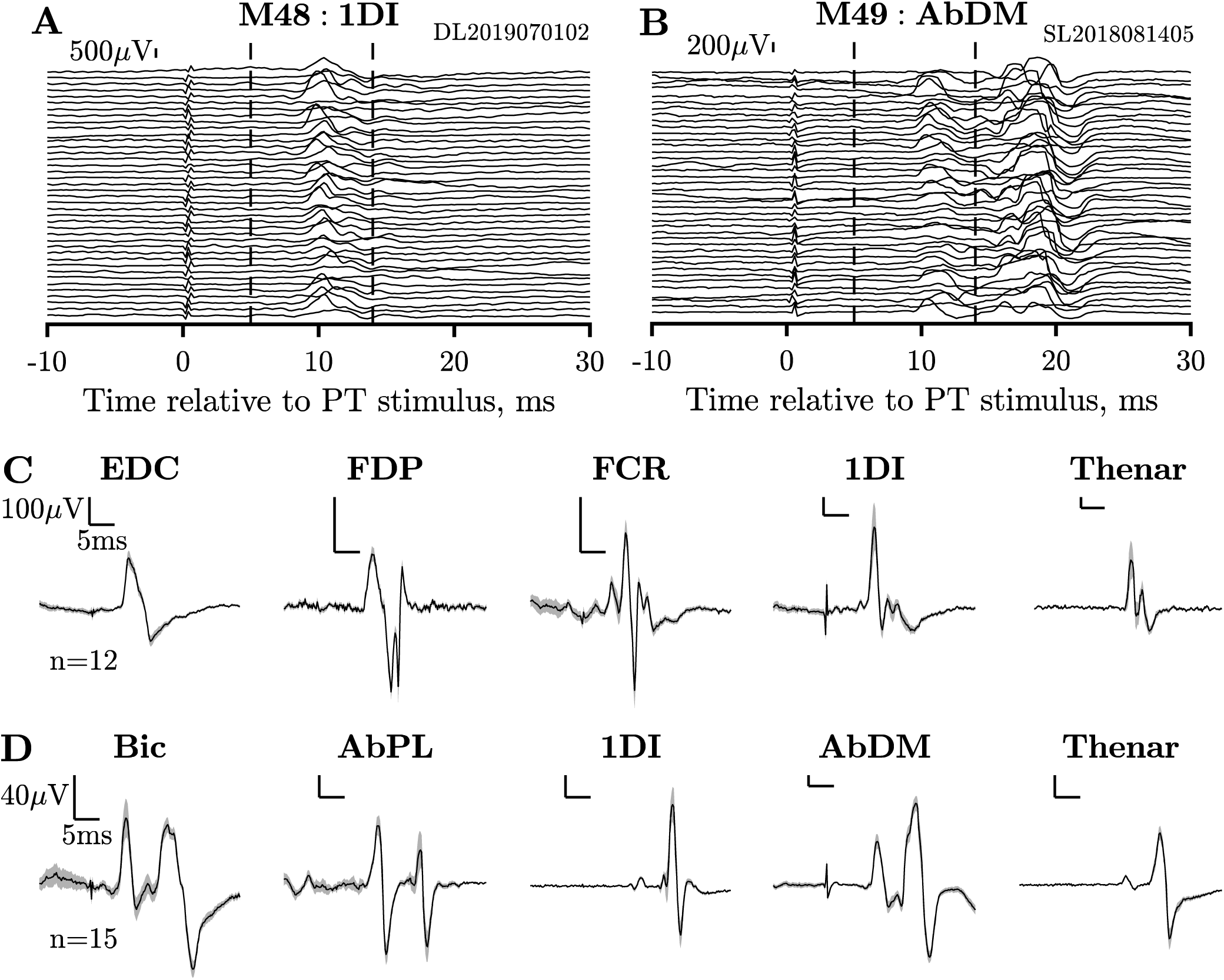
Motor-evoked potentials elicited by PT stimulation. (**A**) Single sweeps of 1DI MEPs recorded in M48 during hand shaping, randomly selected across a single session. Dashed vertical lines denote 5-14ms window used to extract peak-to-peak amplitudes. (**B**) Same as (**A**), but for AbDM in M49. Long-latency responses (∼15-16ms latency) present in the AbDM muscle of M49 during action observation were not included in peak-to-peak analysis window to focus on direct corticospinal effects. (**C**) Responses to stimulation in 5 muscles in M48. Traces show mean+SEM across MEPs during PG hand shaping epoch (300ms leading up to DO) from a single session (note different scales for each muscle). (**D**) Same as (C), but for M49.

### Task-dependent modulation of spinal excitability

To examine the temporal modulation of MEP amplitude during the different conditions, we calculated the mean peak-to-peak amplitude of MEPs within 300ms bins across different stages of the task. Figure 4 shows examples of averaged 1DI MEP responses obtained during *Baseline* and *Hand Shaping* task periods on execution and observation trials for PG and WHG. For PT stimuli during *Baseline*, the resulting MEPs were unsurprisingly comparable across the different task conditions (Fig. 4A). By contrast, during *Hand Shaping* for PG (Fig. 4B), clear differences emerged in the amplitude of MEPs across the different observation conditions; these changes were less pronounced for WHG (Fig. 4C). While MEP amplitudes increased in the Visible Observation and Hidden Grasp conditions, they remained close to baseline in the Hidden Cue&Grasp condition. Execution MEPs, shown on a different scale, were much larger at this point and contaminated by voluntary EMG activity.

**Fig 4.**
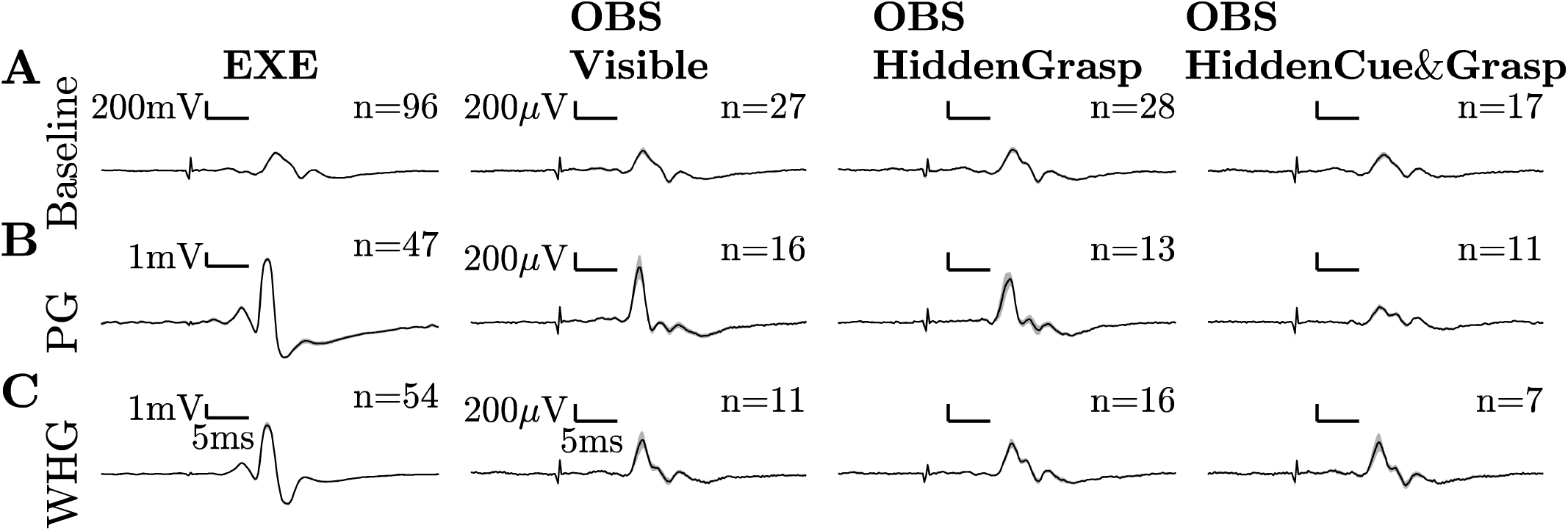
1DI MEPs during different task conditions (one per column) during (**A**) baseline (0-300ms from LCDon) and (**B**) PG *Hand Shaping* (300ms leading up to DO) from a single session in M48. (**C**) Same as (**B**) for WHG. Baseline MEP amplitudes are fairly similar across all conditions. Execution trials are contaminated by voluntary EMG activity and are considerably larger during PG and WHG *Hand Shaping* (note different scale for execution MEPs). Visible Observation and Hidden Grasp trial MEPs increase in amplitude during *Hand Shaping*, particularly for PG, whereas Hidden Cue&Grasp MEPs remain close to baseline.

We next examined the profile of modulation across time, conditions, and sessions. For responses in 1DI of M48 (Fig. 5A), facilitation of spinal excitability during the object presentation period was apparent in all conditions except for Hidden Cue&Grasp observation (green), where LED cues regarding the upcoming trial were not visible, and the MEPs remained close to baseline. MEPs following the imperative Go/NoGo cues then suppressed back to baseline levels or somewhat below, except during execution. Following this, unobstructed observation of PG produced an increase in 1DI MEPs (orange trace), which was particularly prominent during final hand shaping leading up to the experimenter’s grasp. When the grasp was obscured but the monkey had seen the object and Go cues (Hidden Grasp condition), a very similar facilitation profile was observed, with a 50% increase in MEP size in the lead up to grasp. During the Hidden Cue&Grasp condition, on the other hand, when the grasp was obscured and the monkey did not know which object was being grasped, MEPs did not modulate to the same extent. Observation of WHG also showed some facilitation effects around the time of the grasp, but these were weaker than for PG, particularly during the Hidden Grasp condition. During execution, 1DI MEPs were substantially increased from the lead-up to HPR until the end of the trial, reflecting the corresponding increase in spinal excitability during the monkey’s own movements. We observed some qualitatively similar modulation in the AbDM muscle of M49, with an increase in excitability during the observation of hand shaping and grasp for PG, although these effects were generally weaker (Fig. 5B).

**Fig 5.**
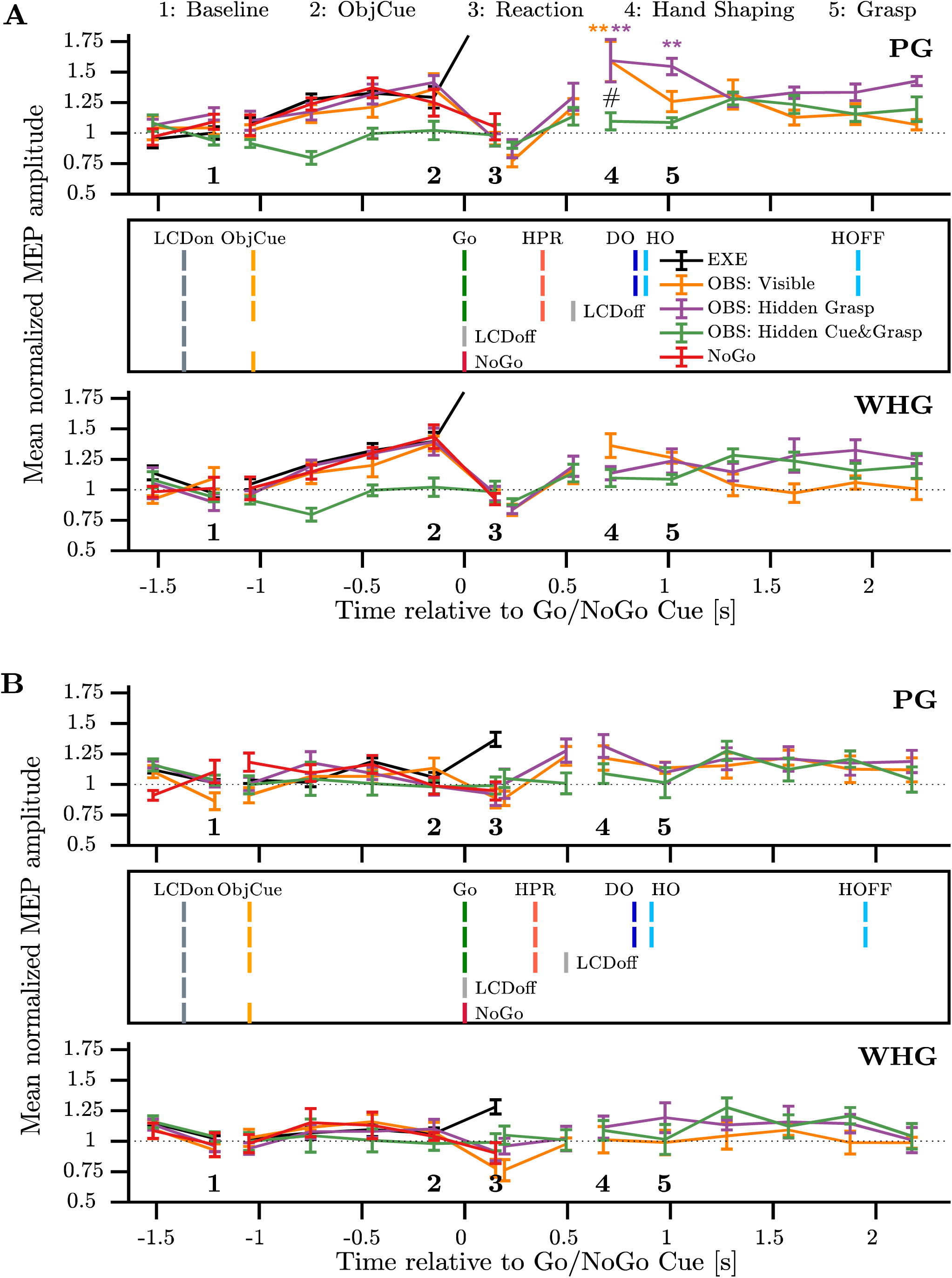
(**A**) Normalized MEP amplitude in 1DI of M48, averaged within 300ms bins across the trial for PG (top) and WHG (bottom), and then across sessions (n=5). Points represent mean±SEM across sessions, with values normalized to mean across conditions in *Baseline* epoch (marked time point 1) within session, before averaging. Execution data is cut off on y-axis to better show modulation during other conditions. Vertical markers in centre show average event times across sessions for five conditions, as indicated by legend. conventions for events as in Fig. 2, with additional markers for the object cue (orange), and time that the screen became opaque on hidden observation trial types Marked time points 1-5 mark bins used for planned comparisons of MEP modulation (*Baseline*, *ObjCue*, *Reaction*, *Hand Shaping*, and *Grasp*). Orange and purple ** indicate significant differences (p<0.005) relative to baseline and # indicates significant (p<0.01) difference across Visible Observation and Hidden Cue&Grasp conditions during Hand Shaping (collapsed across objects). (**B**) Same as (A), but for M49.

To assess quantitatively the modulation of MEP size during action observation, we compared MEP amplitudes across five salient epochs in the observation trial sequence: Baseline, Object Cue, Reaction, *Hand shaping* and Grasp (2-way ANOVA, see Methods). In M48, there was a main effect of EPOCH during Visible Observation and Hidden Grasp conditions in 1DI (both p < 0.0001), and thenar (p < 0.0001 and p = 0.0068, respectively). Post-hoc comparisons across epochs revealed that 1DI MEPs in these two observation conditions were larger during *Hand shaping* relative to *Baseline* (35-40%, both p < 0.005, indicated by orange and purple asterisks on Fig 5A) and *Reaction* (50-65%, both p < 0.0003) epochs (Fig. 5A). The Hidden Grasp condition also showed an interaction of epoch and object (p = 0.045), and 1DI MEPs during hand shaping for PG were significantly larger than those during WHG (p = 0.021). Visible Observation and Hidden Grasp MEPs during the ObjCue epoch were also facilitated relative to those during *Baseline* and *Reaction* in 1DI (all p < 0.05).

In M49, short-latency responses in the 1DI muscle of M49 were generally weak, but more apparent in AbDM (Fig. 3B,D). Nonetheless, a 2-way ANOVA (EPOCH x OBJECT) revealed significant main effects of EPOCH during Visible Observation and Hidden Grasp conditions in 1DI (F_4,40_ > 2.9, p < 0.05), Thenar (F_4,40_ = 4.9, p < 0.003), and AbDM (F_4,40_ = 4.3, p < 0.006). In all three muscles, MEPs during *Hand Shaping* in Visible and Hidden Grasp conditions were 20-35% larger than those in the *Reaction* epoch (all p < 0.03), although modulation relative to Baseline was limited. Across both monkeys and all three muscles, the Hidden Cue&Grasp condition showed no modulation across epochs (all p > 0.3).

Given these results, we next assessed whether visual or predicted information about the observed grasp affected spinal excitability, by comparing directly across the three observation conditions within the *Hand Shaping* epoch (2-way ANOVA). In 1DI of M48, this revealed a significant main effect of condition (F_2,24_ = 5.70, p = 0.009) and object F_1,24_ = 5.84, p =0.024), since PG MEPs were larger than those during WHG. During *hand shaping*, Visible and Hidden Grasp MEPs were not different from each other across objects (p = 0.61), but were larger than those in the Hidden Cue&Grasp condition, by almost 50% in the case of PG (Visible Obs: p = 0.0085, Hidden Grasp: p = 0.07, post-hoc tests collapsed across objects). In the thenar muscle, there was also a main effect of condition (F_2,24_ = 6.38, p = 0.006), and MEPs in Visible and Hidden Grasp conditions were not different to each other (p = 0.53), but again larger than those during Hidden Cue&Grasp (p = 0.0052 and p = 0.06 respectively, post-hoc test collapsed across objects). MEPs in M49 showed no significant differences between observation conditions.

Finally, to assess evidence of suppression associated with action observation and action withholding, we compared MEP amplitudes in the two bins before and after the Go/NoGo cue (*ObjCue* and *Reaction*) across three conditions (Execution, Visible Observation, and NoGo), and two objects (Fig. 6, 3-way ANOVA). All muscles showed significant Condition x Epoch interactions (all p < 0.0001). There was no difference between any of the three conditions during the *ObjCue* period (all p > 0.9). During the Reaction period, which was before any movement (Supplementary Table 1), execution MEPs were consistently larger than Observation and NoGo MEPs, and larger than *ObjCue* MEPs (all p < 0.0001). Observation MEPs showed a 15-30% suppression across monkeys and hand muscles during the Reaction interval relative to *ObjCue*, which was frequently significant (p < 0.05). NoGo MEPs tended to show a similar suppression of 10-30%, and observation and NoGo MEP amplitudes during Reaction were comparable (all p > 0.7).

**Fig 6.**
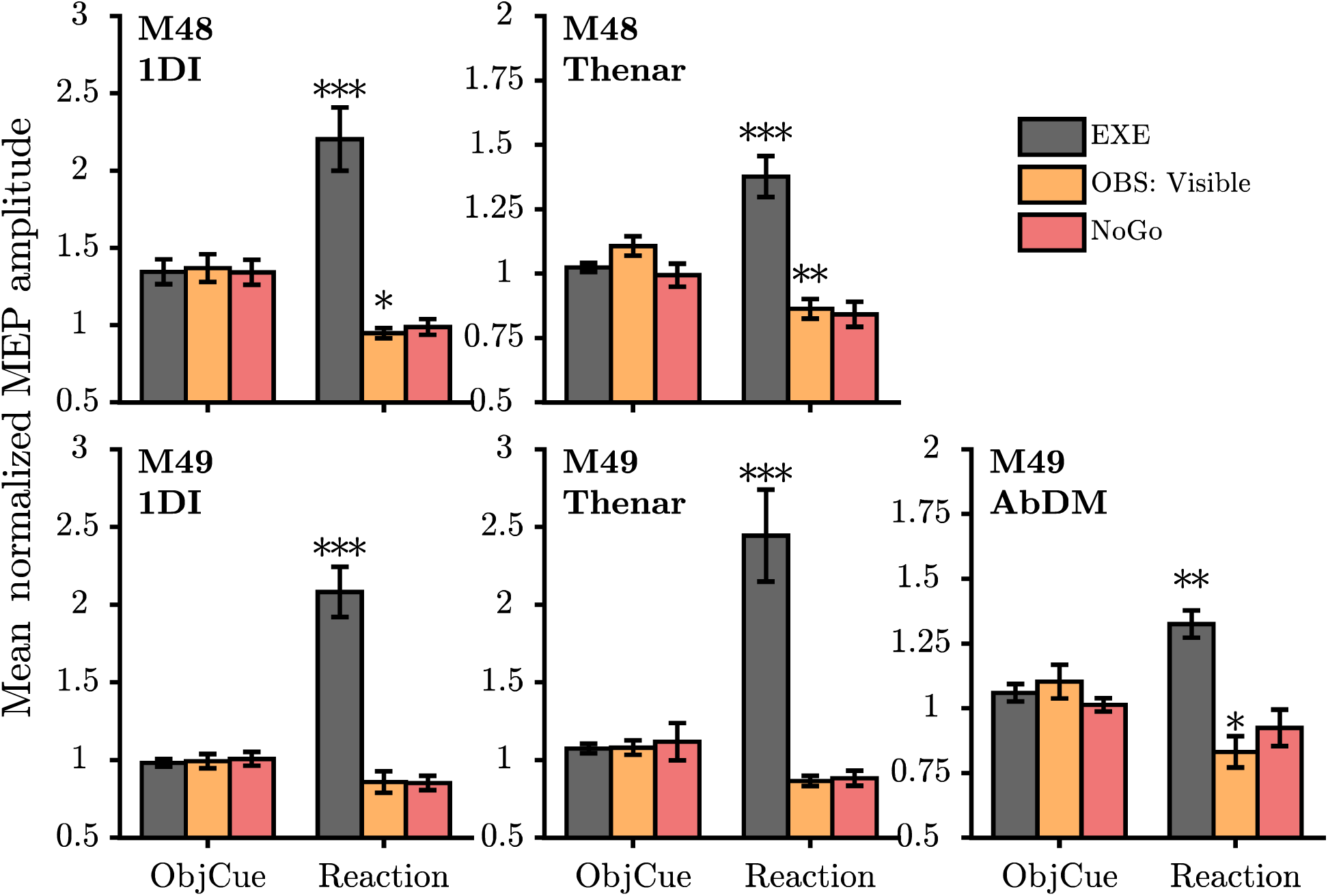
Normalized MEP amplitudes (mean±SEM) in intrinsic hand muscles around the imperative stimulus. Panels show normalized amplitudes across sessions for *ObjCue* and *Reaction* timepoints during Execution, Observation, and NoGo, averaged across objects. Top row: M48, lower row: M49 (n=5 sessions per monkey). Execution MEP amplitude typically increased once movement initiation processes had begun (*Reaction*), whereas Observation and NoGo MEPs during this period were suppressed relative to *ObjCue*. * p < 0.05, ** p < 0.005, *** p < 0.0001, significant difference compared to relevant *ObjCue*. Observation and NoGo were significantly call cases (p < 0.0001), but not from each other (all p > 0.7)

## DISCUSSION

During action observation in macaques, there is modulation in the activity of corticospinal outputs from both premotor and primary motor cortex (Kraskov et al. 2009; Vigneswaran et al. 2013). Similar inferences have been made from human TMS studies probing corticospinal excitability (Fadiga et al. 1995; Gangitano et al. 2001; Montagna et al. 2005; Urgesi et al. 2006; Cattaneo et al. 2009; Bunday et al. 2016), yet these changes do not result in overt movement in the observer. Although spinal circuitry represents the final common path for the neural signals generating or suppressing muscle activity and movement, our understanding of the influence of action observation on this circuitry remains limited.

### Observation of grasp produces facilitation at the spinal level

During the observation of grasp, we found facilitation in the 1DI muscle (Fig. 4 and 5A), consistent with previous results reporting sub-threshold modulation in human corticospinal excitability during action observation (Fadiga et al. 1995; Gangitano et al. 2001; Montagna et al. 2005; Urgesi et al. 2006; Cattaneo et al. 2009; Bunday et al. 2016). In monkeys, there is an overall disfacilitation of corticospinal outputs during action observation (Kraskov et al. 2009; Vigneswaran et al. 2013), but the net effects of this disfacilitation at the spinal level were not tested directly, since the spinal targets of these corticospinal mirror neurons was largely unknown. The results presented here demonstrate that grasp-related mirror activity in descending pathways can produce sub-threshold increases in net excitability within spinal circuitry. This facilitation was most prominent during Hand Shaping before the grasp, and just after the grasp itself. Previous findings in humans suggest that corticospinal excitability modulations peak at the time of observed maximal hand aperture (Gangitano et al. 2001), while dynamic stages of the action are ongoing (Urgesi et al. 2006, 2010), or at the time when grasp is achieved (Gueugneau et al. 2015), and the peak of the observation response in macaque mirror neurons often occurs prior to, or around, the grasp (Vigneswaran et al. 2013). Facilitation in the 1DI muscle was more pronounced for observation of PG rather than WHG (Fig. 5A), although this effect was relatively weak. Grasp and muscle-specific changes in corticospinal excitability occur during action execution (Lemon et al. 1995; Davare et al. 2008, 2009), and action observation (Catmur et al. 2007; Sartori et al. 2012; McCabe et al. 2015; Bunday et al. 2016), and at the cortical level, many macaque mirror neurons show congruence between the executed and observed action most effective in activating them (Gallese et al. 1996). The index finger and 1DI muscle are important for PG execution (Muir and Lemon 1983; Bennett and Lemon 1996; Quallo et al. 2012), and 1DI MEPs are modulated during execution in a time-locked, grasp-specific manner (Davare et al. 2008, 2009). The relatively larger modulation of spinal excitability during PG in the 1DI suggests that action observation activates some of the same spinal circuitry which is used when the monkeys perform the grasp themselves. However, the evidence for specificity of corticospinal excitability modulation during action observation is mixed (Naish et al. 2014), and congruence in cortical mirror neurons is often broad (Gallese et al. 1996). Additionally, the PG and WHG performed by the monkeys in the present task were complex, multi-phasic actions. As such, recording of muscle activity during execution verified that the modulation during observation was complementary to the activation profile of the muscles during execution, demonstrating greater 1DI activation during PG execution than during WHG (Fig. 2A and B). Facilitatory effects during observation of grasp were generally weaker in M49, with the AbDM muscle showing modest facilitation during Hand Shaping relative to Reaction, but not compared to the Baseline (Fig. 5B). One possible explanation for the weaker effect in this monkey may have been the small amplitude of short-latency responses, with several muscles showing late responses with latencies of 15-16ms, i.e. at least 5ms longer than the short latency MEPs (Fig. 3B and D). These long-latency effects were present during action observation, when ongoing EMG activity was absent, and disappeared from MEPs evoked during action execution. They were probably mediated by pathways other than the fast CM component of the corticospinal tract, which might include reticulospinal pathways (Riddle et al. 2009) and propriospinal pathways (Isa et al. 2006; Isa 2019) activated by corticospinal collaterals. However, the additional latency suggests rather indirect, oligosynaptic actions.

A further factor potentially influencing grasp-related facilitation effects during action observation is the behavioural strategy adopted by the two animals, as reflected in their eye-movement patterns (Fig. 2C). Monkeys were allowed to gaze freely during all conditions, which we believe provides a more ethological way to study action observation. M49 consistently diverted gaze from the target object shortly after the Go/NoGo cue which indicated that no response was required, and showed a small tendency to return around the time of the grasp, possibly reacting to the experimenter’s movement. On the other hand, M48 more often maintained gaze on the target object after the experimenter’s movement had begun. A previous study suggested that gaze behaviour during action observation can modulate mirror neuron activity (Maranesi et al. 2013), although almost half of recorded PMv mirror neurons in that study did show gaze-independent modulation during observation. A recent TMS study using simple, single finger movements found that gaze fixation at the point of movement facilitated MEPs relative to free gaze (D’Innocenzo et al. 2017). Given the more complex movements used in the present task, and trial-to-trial variability of free gaze behaviour, further interpretation of the relationship between eye movements and excitability changes in motor pathways during naturalistic action observation, if present, would require simultaneous recordings. From the current data, we speculate that M48 spent more time collecting information about the forthcoming action, presumably for updating of an internal model for better prediction of the upcoming observed grasping action (Kilner et al. 2007).

### Differences in available visual information modulate the response to action observation

A secondary aim of this study was to probe the spinal response during action observation when the amount of visual information available to the monkey was altered. Previously, F5 mirror neurons have been shown to continue to modulate their response even when the grasp was obscured (Umiltá et al. 2001; Kraskov et al. 2009), or only auditory cues are available (Kohler et al. 2002), suggesting that mirror activity is not simply a passive visual response, but an unfolding of the internal representation of the action, reflecting a prediction about the action goal (Kilner et al. 2007). Here we found spinal facilitation was still present in the 1DI of M48 when the view of the grasp was obscured, but the type of grasp was known to the monkey i.e. perfectly predictable. This facilitation showed the same pattern as seen during visible observation (Fig. 5A), with a preference for PG, in which the 1DI muscle is intimately involved (Fig. 2A) (Muir and Lemon 1983; Maier et al. 1993; Quallo et al. 2012). Importantly, the pattern of eye movements in the Hidden Grasp, but not Hidden Cue&Grasp condition, was similar to the Visible Observation condition, suggesting that the monkeys frequently anticipated the target object even though it was not visible, but only if they had seen the object cue. Interestingly, the modulation in the Hidden Grasp condition had a tendency to remain for longer than that seen in the Visible Observation condition. In this latter condition, the monkeys can accurately predict the end stage of the grasp, which may lead to an earlier attenuation of excitability changes (Urgesi et al. 2006, 2010). This persistence of action observation modulation in the absence of direct visual input suggests that, even at the spinal level, the internal representation of an upcoming action may be sufficient to modulate excitability. Previous evidence for comparable changes in corticospinal excitability during hidden action observation is limited, with changes relative to full vision observation and a resting baseline either weak or not tested (Villiger et al. 2011; Valchev et al. 2015). In our task, information about the general trial structure (via hold window and reward sounds) was available to the monkey on both visible and hidden trial types, and auditory cues have been shown to elicit mirror responses (Kohler et al. 2002; Alaerts et al. 2009). These sound cues were not grasp-specific, but may have been sufficient to trigger a more general mirror response in the well-trained monkeys, accounting for some of the non-specific facilitation later in the trial (Fig. 5A).

### Suppression of excitability during suppression of movement

Alongside the grasp-related facilitation during action observation, we were also able to assess spinal excitability at the stage prior to reaching and grasping, at the time when the monkey had to initiate the movement on execution trials, or remain still on observation trials. In both monkeys, we found evidence for suppression of MEPs during this period (Fig. 6) which was comparable in amplitude to when they explicitly suppressed their movement after being cued with a NoGo signal. This suppression following the Go/NoGo cue is consistent with the notion that action observation implicitly requires movement suppression (Kraskov et al. 2009, 2014; Vigneswaran et al. 2013), and suggests that the neural substrate for this suppression overlaps with that involved in the explicit suppression of movement. This finding contrasts with a previous study, which found many mirror neurons which also responded to an observation-NoGo condition, but not when the monkey suppressed its own movement (Bonini et al. 2014). The different findings in our experiment may be due to the use of an interleaved task design and relative timing of the NoGo cue. The monkeys had to decide at matched time points on a trial-by-trial basis whether to generate or suppress movement, which is different from the block design used by Bonini et al. (2014), where it was clear from the outset of all trials within a block whether the monkey would be executing or observing actions. The Go/NoGo cue was then provided as an auditory cue prior to object presentation, meaning that, unlike in our study, the action or inaction could be predicted in advance (Maranesi et al. 2014).

During the ObjCue period preceding the Go/NoGo cue, we observed a consistent increase in excitability across all conditions in which the object cue was provided in M48, with relatively little change in M49. Premotor and motor cortex are known to show anticipatory activity prior to intended movements (Tanji and Evarts 1976; Weinrich and Wise 1982; Alexander and Crutcher 1990; Riehle and Requin 1993; Churchland et al. 2006) and preparation-related changes also occur at the spinal level (Prut and Fetz 1999; Fetz et al. 2002). Suppression of corticospinal excitability during movement preparation has been hypothesised to have important roles in response selection and impulse control (Hasbroucq et al. 1999; Duque and Ivry 2009; Duque et al. 2010; Greenhouse et al. 2015; Lebon et al. 2016). In premotor area F5, a substantial proportion of neurons are active during object presentation, in a manner reflecting upcoming grasp (Murata et al. 1997; Raos et al. 2006; Umiltá et al. 2007). In our task, the non-cued object was still present within the monkey’s field of view for the duration of the trial, and the absence of object specificity during this pre-movement stage suggests this response is distinct from grasp-specific facilitation reported in F5 neurons during object presentation (Raos et al. 2006). We therefore consider it more likely that the different levels of modulation during *ObjCue* in the two monkeys arise due to qualitatively different strategies. Eye movement recordings suggest that M48 was more attentive during action observation than M49 (Fig. 2C), and in extracellular recordings of PTNs in M1 in the same monkeys (unpublished observations), we generally found higher proportions, and greater modulation, of mirror neurons in M48. Along with possible inhibitory mechanisms at cortical and sub-cortical levels, attentional and motivational factors of the specific monkey may influence activity in cortical and spinal motor circuitry during action observation.

## Conclusions

Here, we used direct stimulation of the pyramidal tract to probe the modulation of spinal excitability during preparation, observation, and explicit suppression of reach-to-grasp actions. The MEPs elicited by PT stimulation reflect post-synaptic mechanisms in α-motoneurons. PT stimulation therefore offers a useful and unique method for probing excitability of spinal circuitry controlling the hand in the awake, behaving animal.

Our results confirm that the motoneuron pools innervating the hand undergo sub-threshold modulations during action observation. We found an increase in excitability at the spinal level around the observation of hand shaping and grasp, particularly in the 1DI muscle during PG, suggesting that the same spinal circuits which are recruited during PG performance are modulated when the same action is observed. This excitability was still apparent when the grasp was obscured but predictable, indicating that predictive activity of cortical mirror neurons can reach and excite motoneurons in the spinal cord. This increase in excitability was preceded by a relative suppression in the lead-up to the observation of movement. This was comparable to the suppression when the monkey withheld movement, and may support the suppression of movement during action observation. Our previous findings showed that descending activity from PTNs in PMv, and particularly in M1, is attenuated during observation, and we inferred from those results that hand motoneurons would receive less excitation than during execution. Here we used a more direct assessment of spinal motoneuron excitability and the relatively modest effects, and balance of excitation and inhibition, provide further evidence that the net effects of motor cortical output during the different stages of action observation must be limited such that spinal motoneuron pools remain sub-threshold to movement.

Future studies might consider more direct measures of spinal modulation during action observation, including single motor unit responses to PT stimulation, as well as extracellular recording from spinal interneurons. These could shed light on the excitatory and inhibitory effects of cortical mirror neuron activity on the spinal cord machinery. More generally, this could yield further insights in to the physiological mechanisms underpinning the generation and suppression of grasping movements.

## Contributions

S.J.J. performed experiments, analysed data, prepared figures, and drafted the manuscript. S.J.J., R.N.L. and A.K. designed research, discussed results, revised the manuscript, and approved the final version. A.K. supervised the project.

## Acknowledgements

The authors thank Tabatha Lawton, Dominika Klisko and Adam Keeler for help with recordings, and Spencer Neal, Jonathon Henton, Chris Seers, & Martin Lawton for technical assistance, and Dr Marco Davare for useful discussions of this project.

---

**Supplementary Table 1:** Behaviour

|  | M48 |  | M49 |  |
| --- | --- | --- | --- | --- |
|  | Monkey | Human | Monkey | Human |
| # Trials<br>(PG,WHG) | 734<br>(346, 388) | 258<br>(127, 131) | 820<br>(417,403) | 258<br>(127,131) |
| Reaction Time | 0.31 ± 0.07 | 0.42 ± 0.06 | 0.27 ± 0.03 | 0.40 ± 0.07 |
| Movement Time (PG) | 0.42 ± 0.11 | 0.51 ± 0.07 | 0.39 ± 0.05 | 0.50 ± 0.05 |
| Movement Time (WHG) | 0.33 ± 0.07 | 0.45 ± 0.10 | 0.33 ± 0.03 | 0.51 ± 0.05 |

Reaction time (across objects), and movement time for each object displayed as mean±S.D. in seconds, across trials from all 5 sessions in each monkey. Human trials include Visible Observation trials only.

**Supplementary Table 2:**
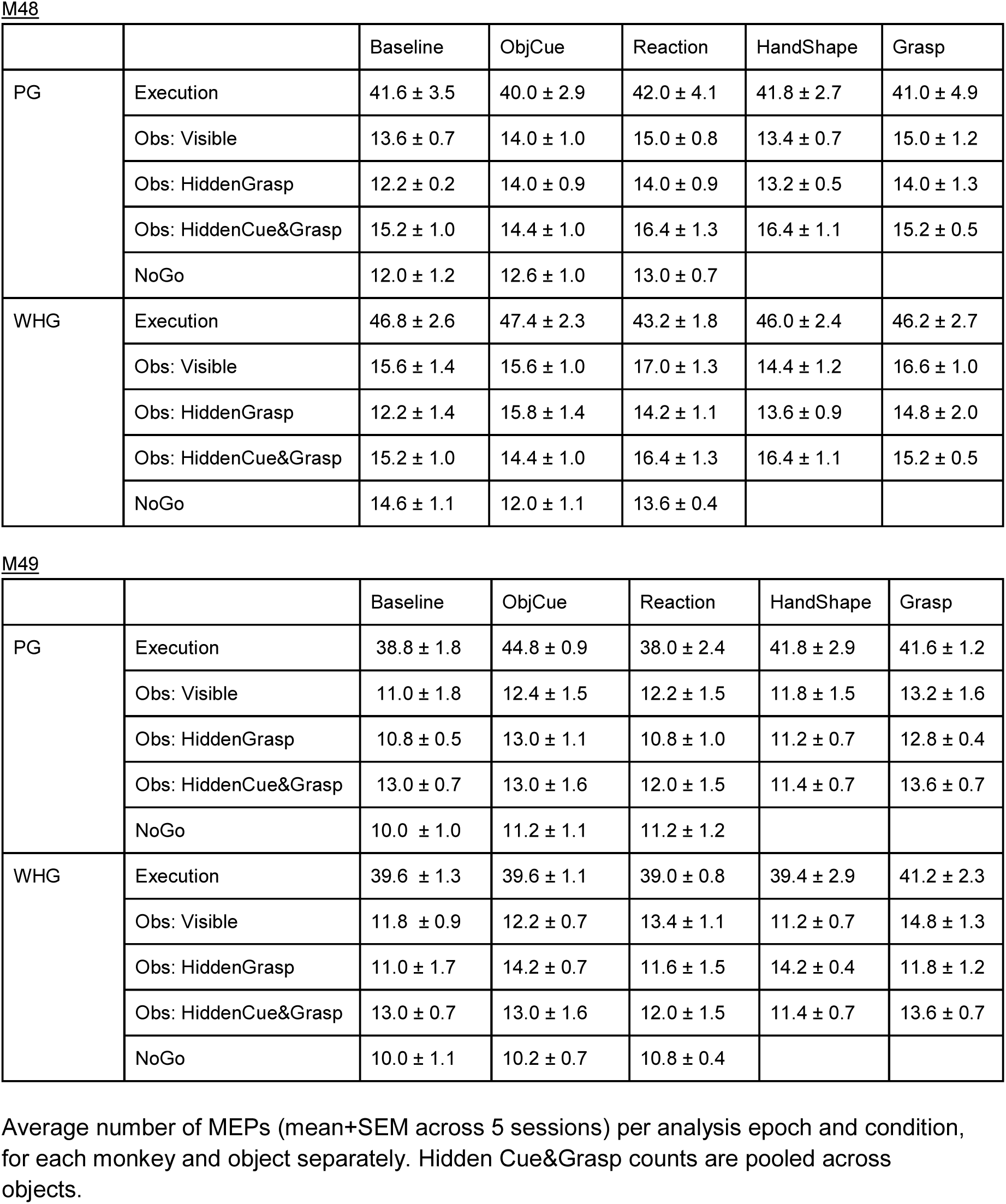
Number of MEPs used for analysis

## REFERENCES

1. Alaerts K, Swinnen SP, Wenderoth N. 2009. Interaction of sound and sight during action perception: Evidence for shared modality-dependent action representations. Neuropsychologia. 47:2593–2599.

2. Alexander GE, Crutcher MD. 1990. Preparation for movement: neural representations of intended direction in three motor areas of the monkey. J Neurophysiol. 64:133–150.

3. Baldissera F, Cavallari P, Craighero L, Fadiga L. 2001. Modulation of spinal excitability during observation of hand actions in humans. Eur J Neurosci. 13:190–194.

4. Bennett KM, Lemon RN. 1996. Corticomotoneuronal contribution to the fractionation of muscle activity during precision grip in the monkey. J Neurophysiol. 75:1826–1842.

5. Bonini L, Maranesi M, Livi A, Fogassi L, Rizzolatti G. 2014. Ventral premotor neurons encoding representations of action during self and others’ inaction. Curr Biol. 24:1611– 1614.

6. Borra E, Belmalih A, Gerbella M, Rozzi S, Luppino G. 2010. Projections of the hand field of the macaque ventral premotor area F5 to the brainstem and spinal cord. J Comp Neurol. 518:2570–2591.

7. Borroni P, Montagna M, Cerri G, Baldissera F. 2005. Cyclic time course of motor excitability modulation during the observation of a cyclic hand movement. Brain Res. 1065:115– 124.

8. Bunday KL, Lemon RN, Kilner JM, Davare M, Orban GA. 2016. Grasp-specific motor resonance is influenced by the visibility of the observed actor. Cortex. 84:43–54.

9. Catmur C, Walsh V, Heyes C. 2007. Sensorimotor Learning Configures the Human Mirror System. Curr Biol. 17:1527–1531.

10. Cattaneo L, Caruana F, Jezzini A, Rizzolatti G. 2009. Representation of Goal and Movements without Overt Motor Behavior in the Human Motor Cortex: A Transcranial Magnetic Stimulation Study. J Neurosci. 29:11134–11138.

11. Cerri G, Shimazu H, Maier MA, Lemon RN. 2003. Facilitation From Ventral Premotor Cortex of Primary Motor Cortex Outputs to Macaque Hand Muscles. J Neurophysiol. 90:832–842.

12. Churchland MM, Yu BM, Ryu SI, Santhanam G, Shenoy K V. 2006. Neural Variability in Premotor Cortex Provides a Signature of Motor Preparation. J Neurosci. 26:3697–3712.

13. D’Innocenzo G, Gonzalez CC, Nowicky A V., Williams AM, Bishop DT. 2017. Motor resonance during action observation is gaze-contingent: A TMS study. Neuropsychologia. 103:77–86.

14. Davare M, Lemon R, Olivier E. 2008. Selective modulation of interactions between ventral premotor cortex and primary motor cortex during precision grasping in humans. J Physiol. 586:2735–2742.

15. Davare M, Montague K, Olivier E, Rothwell JC, Lemon RN. 2009. Ventral premotor to primary motor cortical interactions during object-driven grasp in humans. Cortex. 45:1050–1057.

16. Dum RP, Strick PL. 2005. Frontal Lobe Inputs to the Digit Representations of the Motor Areas on the Lateral Surface of the Hemisphere. J Neurosci. 25:1375–1386.

17. Duque J, Ivry RB. 2009. Role of corticospinal suppression during motor preparation. Cereb Cortex. 19:2013–2024.

18. Duque J, Lew D, Mazzocchio R, Olivier E, Ivry RB. 2010. Evidence for Two Concurrent Inhibitory Mechanisms during Response Preparation. J Neurosci. 30:3793–3802.

19. Fadiga L, Fogassi L, Pavesi G, Rizzolatti G. 1995. Motor facilitation during action observation: a magnetic stimulation study. J Neurophysiol. 73:2608–2611.

20. Fetz EE, Perlmutter SI, Prut Y, Seki K, Votaw S. 2002. Roles of primate spinal interneurons in preparation and execution of voluntary hand movement. Brain Res Rev. 40:53–65.

21. Gallese V, Fadiga L, Fogassi L, Rizzolatti G. 1996. Action recognition in the premotor cortex. Brain. 119:593–609.

22. Gangitano M, Mottaghy FM, Pascual-Leone A. 2001. Phase-specific modulation of cortical motor output during movement observation. Neuroreport. 12:1489–1492.

23. Godschalk M, Lemon RN, Kuypers HGJM, Ronday HK. 1984. Cortical afferents and efferents of monkey postarcuate area: an anatomical and electrophysiological study. Exp Brain Res. 56:410–424.

24. Greenhouse I, Sias A, Labruna L, Ivry RB. 2015. Nonspecific Inhibition of the Motor System during Response Preparation. J Neurosci. 35:10675–10684.

25. Gueugneau N, Mc Cabe SI, Villalta JI, Grafton ST, Della-Maggiore V. 2015. Direct mapping rather than motor prediction subserves modulation of corticospinal excitability during observation of actions in real time. J Neurophysiol. 113:3700–3707.

26. Hannah R, Rocchi L, Rothwell JC. 2018. Observing without acting: A balance of excitation and suppression in the human corticospinal pathway? Front Neurosci. 12:1–10.

27. Hasbroucq T, Kaneko H, Akamatsu M, Possamaï CA. 1999. The time-course of preparatory spinal and cortico-spinal inhibition: An H-reflex and transcranial magnetic stimulation study in man. Exp Brain Res. 124:33–41.

28. He SQ, Dum RP, Strick PL. 1993. Topographic organization of corticospinal projections from the frontal lobe: Motor areas on the lateral surface of the hemisphere. J Neurosci. 13:952–980.

29. Isa T. 2019. Dexterous Hand Movements and Their Recovery After Central Nervous System Injury. Annu Rev Neurosci. 42:315–335.

30. Isa T, Ohki Y, Seki K, Alstermark B. 2006. Properties of Propriospinal Neurons in the C3-C4 Segments Mediating Disynaptic Pyramidal Excitation to Forelimb Motoneurons in the Macaque Monkey. J Neurophysiol. 95:3674–3685.

31. Jackson A, Baker SN, Fetz EE. 2006. Tests for presynaptic modulation of corticospinal terminals from peripheral afferents and pyramidal tract in the macaque. J Physiol. 573:107–120.

32. Kilner JM, Friston KJ, Frith CD. 2007. Predictive coding: An account of the mirror neuron system. Cogn Process. 8:159–166.

33. Kilner JM, Lemon RN. 2013. What we know currently about mirror neurons. Curr Biol. 23:1057–1062.

34. Knikou M. 2008. The H-reflex as a probe: Pathways and pitfalls. J Neurosci Methods. 171:1–12.

35. Kohler E, Keysers C, Umiltá MA, Fogassi L, Gallese V, Rizzolatti G. 2002. Hearing Sounds, Understanding Actions: Action Representation in Mirror Neurons. Science (80-). 297:846–848.

36. Kraskov A, Dancause N, Quallo MM, Shepherd S, Lemon RN. 2009. Corticospinal Neurons in Macaque Ventral Premotor Cortex with Mirror Properties: A Potential Mechanism for Action Suppression? Neuron. 64:922–930.

37. Kraskov A, Philipp R, Waldert S, Vigneswaran G, Quallo MM, Lemon RN. 2014. Corticospinal mirror neurons. Philos Trans R Soc B - Biol Sci. 369:20130174.

38. Kraskov A, Prabhu G, Quallo MM, Lemon RN, Brochier T. 2011. Ventral premotor-motor cortex interactions in the macaque monkey during grasp: Response of single neurons to intracortical microstimulation. J Neurosci. 31:8812–8821.

39. Lebon F, Greenhouse I, Labruna L, Vanderschelden B, Papaxanthis C, Ivry RB. 2016. Influence of Delay Period Duration on Inhibitory Processes for Response Preparation. Cereb Cortex. 26:2461–2470.

40. Lemon RN. 2008. Descending Pathways in Motor Control. Annu Rev Neurosci. 31:195–218.

41. Lemon RN, Johansson RS, Westling G. 1995. Corticospinal Control during Reach, Grasp, and Precision Lift in Man. J Neurosci. 15:6145–6156.

42. Maier MA, Bennett KM, Hepp-Reymond MC, Lemon RN. 1993. Contribution of the monkey corticomotoneuronal system to the control of force in precision grip. J Neurophysiol. 69:772–785.

43. Maier MA, Kirkwood PA, Brochier T, Lemon RN. 2013. Responses of single corticospinal neurons to intracortical stimulation of primary motor and premotor cortex in the anesthetized macaque monkey. J Neurophysiol. 109:2982–2998.

44. Maranesi M, Livi A, Fogassi L, Rizzolatti G, Bonini L. 2014. Mirror Neuron Activation Prior to Action Observation in a Predictable Context. J Neurosci. 34:14827–14832.

45. Maranesi M, Ugolotti Serventi F, Bruni S, Bimbi M, Fogassi L, Bonini L. 2013. Monkey gaze behaviour during action observation and its relationship to mirror neuron activity. Eur J Neurosci. 38:3721–3730.

46. Matelli M, Camarda R, Glickstein M, Rizzolatti G. 1986. Afferent and efferent projections of the inferior area 6 in the macaque monkey. J Comp Neurol. 251:281–298.

47. Mazzocchio R, Rothwell JC, Rossi A. 1995. Distribution of Ia effects onto human hand muscle motoneurones as revealed using an H reflex technique. J Physiol. 489:263–273.

48. McCabe SI, Villalta JI, Saunier G, Grafton ST, Della-Maggiore V. 2015. The Relative Influence of Goal and Kinematics on Corticospinal Excitability Depends on the Information Provided to the Observer. Cereb Cortex. 25:2229–2237.

49. McKiernan BJ, Marcario JK, Karrer JH, Cheney PD. 1998. Corticomotoneuronal Postspike Effects in Shoulder, Elbow, Wrist, Digit, and Intrinsic Hand Muscles During a Reach and Prehension Task. J Neurophysiol. 80:1961–1980.

50. Montagna M, Cerri G, Borroni P, Baldissera F. 2005. Excitability changes in human corticospinal projections to muscles moving hand and fingers while viewing a reaching and grasping action. Eur J Neurosci. 22:1513–1520.

51. Morecraft RJ, Ge J, Stilwell-Morecraft KS, McNeal DW, Pizzimenti MA, Darling WG. 2013. Terminal distribution of the corticospinal projection from the hand/arm region of the primary motor cortex to the cervical enlargement in rhesus monkey. J Comp Neurol. 521:4205–4235.

52. Muakkassa KF, Strick PL. 1979. Frontal lobe inputs to primate motor cortex: evidence for four somatotopically organized “premotor” areas. Brain Res. 177:176–182.

53. Muir RB, Lemon RN. 1983. Corticospinal neurons with a special role in precision grip. Brain Res. 261:312–316.

54. Murata A, Fadiga L, Fogassi L, Gallese V, Raos V, Rizzolatti G. 1997. Object Representation in the Ventral Premotor Cortex (Area F5) of the Monkey. J Neurophysiol. 78:2226–2230.

55. Naish KR, Houston-Price C, Bremner AJ, Holmes NP. 2014. Effects of action observation on corticospinal excitability: Muscle specificity, direction, and timing of the mirror response. Neuropsychologia. 64:331–348.

56. Nakajima K, Maier MA, Kirkwood PA, Lemon RN. 2000. Striking differences in transmission of corticospinal excitation to upper limb motoneurons in two primate species. J Neurophysiol. 84:698–709.

57. Olivier E, Baker SN, Lemon RN, Brochier T, Nakajima K. 2001. Investigation Into Non-Monosynaptic Corticospinal Excitation of Macaque Upper Limb Single Motor Units. J Neurophysiol. 86:1573–1586.

58. Porter R, Lemon RN. 1993. Corticospinal function and voluntary movement. Oxford: Clarendon Press.

59. Prabhu G, Shimazu H, Cerri G, Brochier T, Spinks RL, Maier MA, Lemon RN. 2009. Modulation of primary motor cortex outputs from ventral premotor cortex during visually guided grasp in the macaque monkey. J Physiol. 587:1057–1069.

60. Prut Y, Fetz EE. 1999. Primate spinal interneurons show pre-movement instructed delay activity. Nature. 401:590–594.

61. Quallo MM, Kraskov A, Lemon RN. 2012. The activity of primary motor cortex corticospinal neurons during tool use by macaque monkeys. J Neurosci. 32:17351–17364.

62. Raos V, Umiltá MA, Murata A, Fogassi L, Gallese V, Fogassi L. 2006. Functional Properties of Grasping-Related Neurons in the Ventral Premotor Area F5 of the Macaque Monkey. J Neurophysiol. 95:709–729.

63. Rathelot J-A, Strick PL. 2006. Muscle representation in the macaque motor cortex: An anatomical perspective. Proc Natl Acad Sci USA. 103:8257–8262.

64. Riddle CN, Edgley SA, Baker SN. 2009. Direct and Indirect Connections with Upper Limb Motoneurons from the Primate Reticulospinal Tract. J Neurosci. 29:4993–4999.

65. Riehle A, Requin J. 1993. The predictive value for performance speed of preparatory changes in neuronal activity of the monkey motor and premotor cortex. Behav Brain Res. 53:35–49.

66. Rizzolatti G, Fadiga L, Gallese V, Fogassi L. 1996. Premotor cortex and the recognition of motor actions. Cogn Brain Res. 3:131–141.

67. Sartori L, Bucchioni G, Castiello U. 2012. Motor cortex excitability is tightly coupled to observed movements. Neuropsychologia. 50:2341–2347.

68. Schmidlin E, Brochier T, Maier MA, Kirkwood PA, Lemon RN. 2008. Pronounced Reduction of Digit Motor Responses Evoked from Macaque Ventral Premotor Cortex after Reversible Inactivation of the Primary Motor Cortex Hand Area. J Neurosci. 28:5772–5783.

69. Shimazu H, Maier MA, Cerri G, Kirkwood PA, Lemon RN. 2004. Macaque ventral premotor cortex exerts powerful facilitation of motor cortex outputs to upper limb motoneurons. J Neurosci. 24:1200–1211.

70. Strafella AP, Paus T. 2000. Modulation of cortical excitability during action observation: a transcranial magnetic stimulation study. Neuroreport. 11:2289–2292.

71. Tanji J, Evarts E V. 1976. Anticipatory activity of motor cortex neurons in relation to direction of an intended movement. J Neurophysiol. 39:1062–1068.

72. Umiltá MA, Brochier T, Spinks RL, Lemon RN. 2007. Simultaneous recording of macaque premotor and primary motor cortex neuronal populations reveals different functional contributions to visuomotor grasp. J Neurophysiol. 98:488–501.

73. Umiltá MA, Kohler E, Gallese V, Fogassi L, Fadiga L, Keysers C, Rizzolatti G. 2001. I Know What You Are Doing: A Neurophysiological Study. Neuron. 31:155–165.

74. Urgesi C, Maieron M, Avenanti A, Tidoni E, Fabbro F, Aglioti SM. 2010. Simulating the future of actions in the human corticospinal system. Cereb Cortex. 20:2511–2521.

75. Urgesi C, Moro V, Candidi M, Aglioti SM. 2006. Mapping Implied Body Actions in the Human Motor System. J Neurosci. 26:7942–7949.

76. Valchev N, Zijdewind I, Keysers C, Gazzola V, Avenanti A, Maurits NM. 2015. Weight dependent modulation of motor resonance induced by weight estimation during observation of partially occluded lifting actions. Neuropsychologia. 66.

77. Vigneswaran G, Philipp R, Lemon RN, Kraskov A. 2013. M1 corticospinal mirror neurons and their role in movement suppression during action observation. Curr Biol. 23:236–243.

78. Villiger M, Chandrasekharan S, Welsh TN. 2011. Activity of human motor system during action observation is modulated by object presence. Exp Brain Res. 209:85–93.

79. Weinrich M, Wise SP. 1982. The premotor cortex of the monkey. J Neurosci. 2:1329–1345.

